# Polycyclic Aromatic Hydrocarbon, Heavy Metal, and Derivable Metabolic Water and Energy of Catle Hides Processed by Singeing

**DOI:** 10.1101/2020.03.18.994962

**Authors:** Eze C. Woko, C. O. Ibegbulem, Chinwe S. Alisi

## Abstract

Meat is consumed as source of protein, processed cattle hide popularly known as *Kanda* in southeastern Nigeria is consumed as a substitute for meat. The commercial method of processing this very food delicacy through singeing with scrap tyre, or firewood has posed question marks over the contamination status of the *Kanda*. This study therefore investigated the heavy metal, polycyclic aromatic hydrocarbon (PAHs) content, proximate composition, and also calculated the derivable metabolic water and energy content of *Kanda* singed with scrap tyre, and firewood. Singeing significantly reduced (p < 0.05) the moisture, ash, fibre and carbohydrate contents of the hide while it increased the protein content; increased (p < 0.05) the iron (Fe) concentration of the hide though the concentration of iron detected was far below the average lethal dose allowed in meat and meat products and elevated (p < 0.05) the PAH concentration of the hides. However, the hides singed with firewood showed higher (p < 0.05) concentrations of benzo[a]pyrene. With the exception of naphthalene, 2-methylnaphthalene, acenenaphthalene and indeno [1,2,3-cd] pyrene all the 16 Environmental Protection Agency (EPA) priority PAHs evaluated showed concentration above 0.0001 ppm (0.1 μg/kg) in at least one of the singed samples. Also, Singeing significantly increased (p < 0.05) the derivable metabolic water and energy contents of the cattle hides. In conclusion, while consuming singed cattle hide (*Kanda*) may offer higher energy content and derivable metabolic water, it may in the long term expose the consumer to health hazards associated with heavy metal and PAH contamination.

## 1. Introduction

Food contamination is a worldwide challenge which has been recognized by the World Health Organization (WHO) (Hussain, 2016). In Aba (southeastern Nigeria) singeing with scrap tyre, or firewood have become a major method of removing the hair on the skin of slaughtered cattles and goats (plate 1 and 2). This disturbing method is not unique to Aba alone, but has been observed in several other parts of Africa and is worrying since it can introduce different contaminants into the meat, thereby rendering it unsafe for human consumption (Obiri-Danso *et al*., 2008; Richard *et al*., 2014). It is against this background that this study intends to evaluate the proximate composition, heavy metal, polycyclic aromatic hydrocarbon (PAH) accumulation in scrap tyre-singed hide compared to that singed with fire wood, and an unsigned sample, and also to determine its derivable metabolic water and energy content.

**Plate 1:**
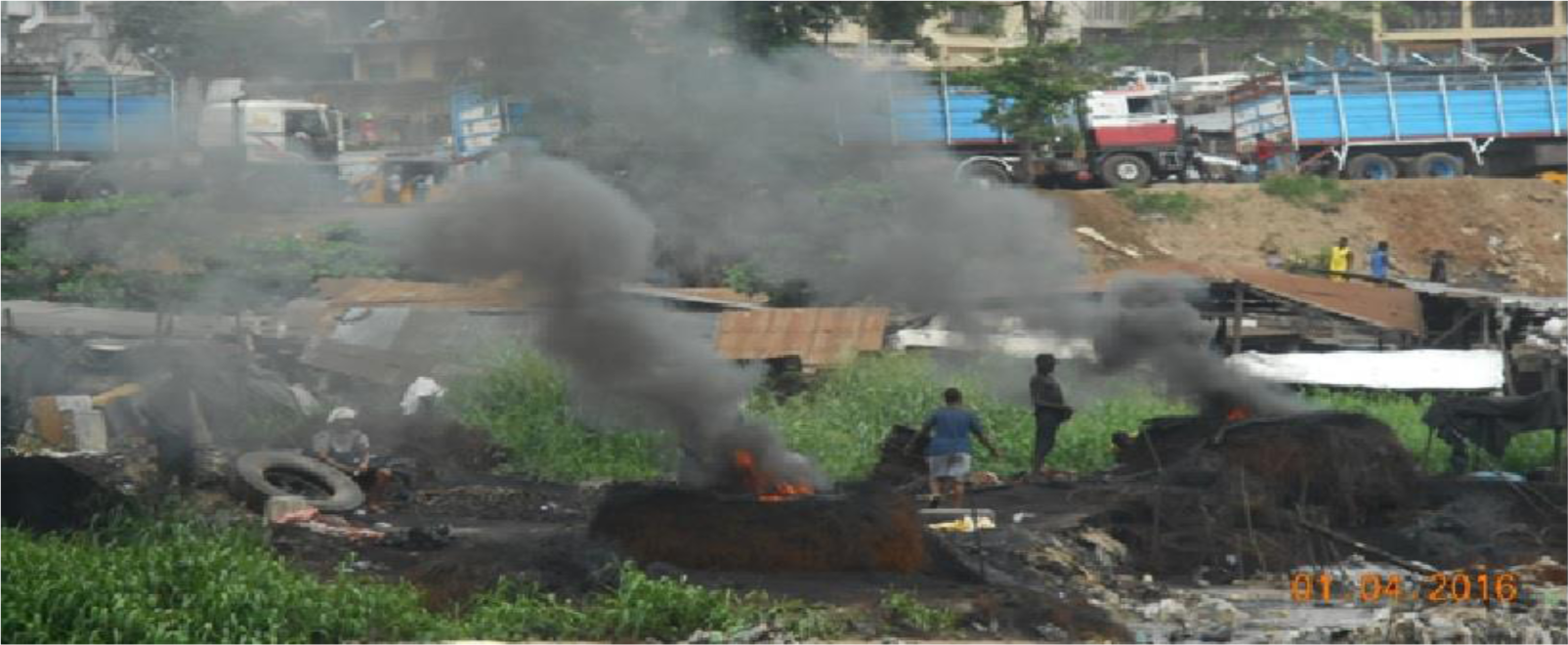
Open burning of scrap tyres for singeing of cattle hides at Ogbor-Hill Aba, Nigeria.

**Plate 2 (A and B):**
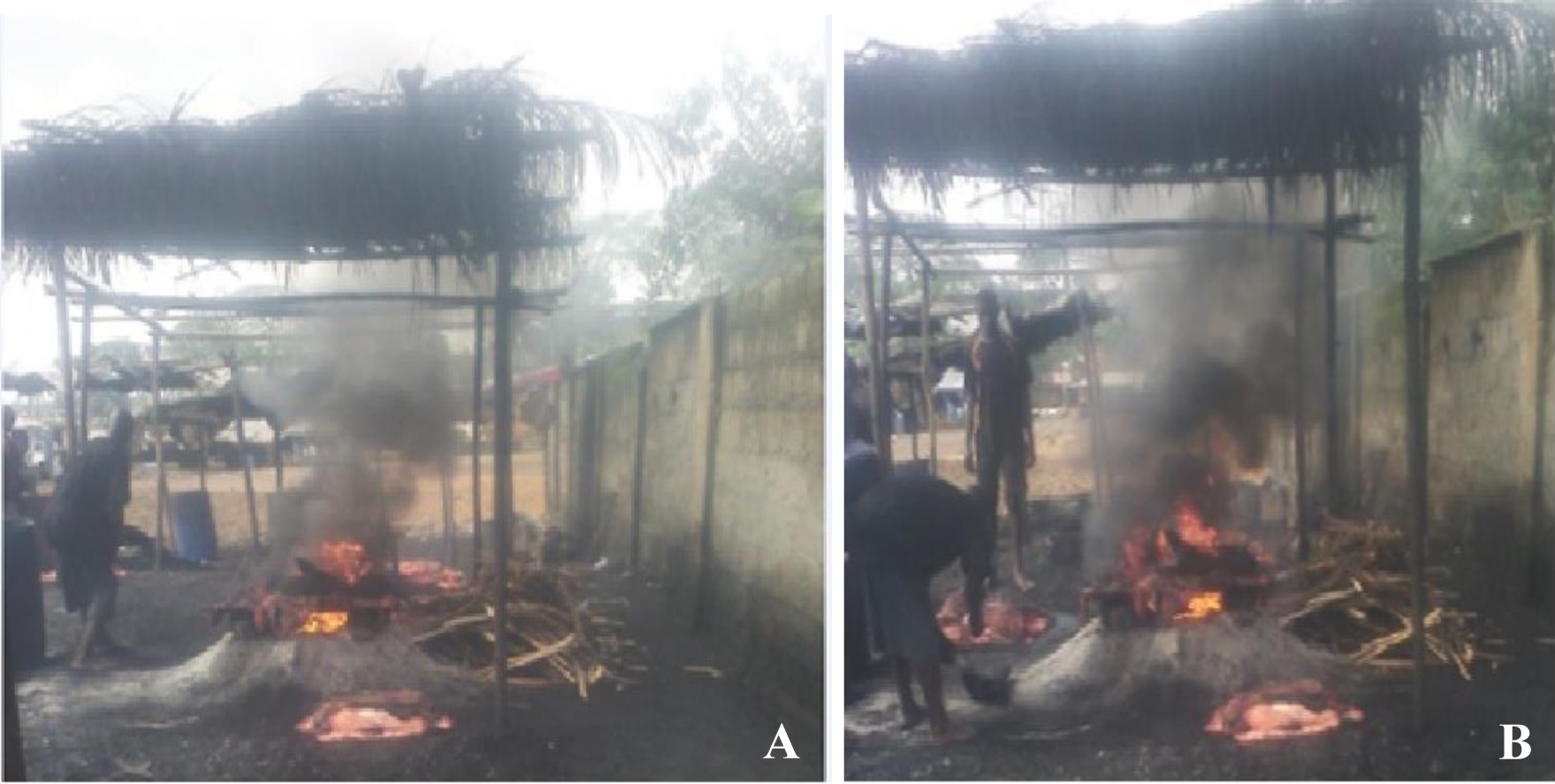
Singeing of cattle hide with firewood at Ogbor-Hill Aba Abattoir, Nigeria.

The risk of meat getting contaminated with heavy metals is a subject of great concern for both food safety and human health because of the toxic nature of these metals at relatively minute concentrations (Santhi *et al*., 2008). Polycyclic aromatic hydrocarbons (PAHs) are a group of environmental contaminants that come from the incomplete combustion of organic matter which over the years have drawn much scrutiny due to their ability to cause cancer (Akpambang *et al*., 2009). When ingested, unmetabolized PAHs can have toxic effects, but the ability of its reactive metabolites such as epoxies and dihydrodiols to bind to cellular proteins and DNA with its resultant biochemical disruptions and cell damage leading to mutations, developmental malformations, tumours and cancer is of major concern (Nisha *et al*., 2015). Generally contamination of meat depends on the nature and age of the animal, place of animal rearing, dietary habits, slaughtering, transportation condition and exposure time (Sabir *et al*., 2003).

## 2. Materials and methods

### 2.1 Procurement and preparation of samples

Freshly slaughtered cattle hides used for this study were purchased from Ogbor Hill Aba slaughter market, Abia state, Nigeria and processed by the author. The hide was then divided into three groups; A, B and C. Sample A was processed by singeing with scrap tyre, the resulting soothe was washed off with clean water. Sample B was processed by singeing with firewood and the resulting soothe was washed off with clean water. Sample C (control) was processed using newly purchased razor blades to scrap off the furs, and then washed with clean water. Each of Samples A, B and C after processing were then further divided into three (3) equal parts and sent to the laboratory for determination of proximate composition, heavy metal and polycyclic aromatic hydrocarbon contents (the groups separated for PAHs analysis were wrapped with aluminum foil to prevent exposure to light).

### 2.2 Determination of proximate composition

Samples of cattle hides processed with scrap tyre, firewood, and control were analyzed for proximate composition according to the method of AOAC (2006). The percentage carbohydrate was conveniently estimated by the FAO difference method (FAO, 2002).

### 2.3 Determination of derivable metabolic water and energy content

The derivable metabolic water content of the samples were calculated on the basis of 0.41 g H_2_O/100 g protein, 0.60 g H_2_O/100 g carbohydrate and 1.07 g H_2_O/100 g fat (lipid) as described by Morrison (1953). While the derivable energy values of each of the samples were calculated using the Atwater factors of 4, 9, 4 in Kcal per 100 g as described by Amadi *et al*. (2004).

### 2.4 Determination of heavy metal content

The heavy metal contents of the samples were determined using the dry-ashing method as described by Aigberua and Tarawou (2017).

### 2.5 Determination of the polycyclic aromatic hydrocarbon content

The polycyclic aromatic hydrocarbon contents of the samples were determined using gas chromatography (GC-MS Spectrometer model 6800) according to the method described by TNRCC (1997).

### 2.6 Statistical analysis

Data generated from proximate, heavy metal and polycyclic aromatic hydrocarbon analysis were evaluated using coefficient of variation statistical method and the means were declared significantly different at 5% (*P* ≤ 0.05).

## 3. Results

The analysis of seven (7) heavy metal compositions of the hides (table 1) revealed only the presence of copper (Cu) and iron (Fe) in all the samples while the other heavy metals were not detected. There was a significant (p<0.05) difference in iron (Fe) when sample A was compared with sample B and sample C, but the iron content of sample B showed no significant (p<0.05) difference when compared with the control (sample C). Sample A and sample B showed significant (p<0.05) difference in copper content when compared, and both samples significantly varied with the control (sample C). The sixteen (16) Environmental Protection Agency (EPA) priority PAHs analysis of the hides (table 2) revealed the presence of all these PAHs at varying concentrations in all the samples. The proximate composition of the hides (table 3) showed that singeing significantly (p<0.05) decreased the moisture, fibre and carbohydrate contents of the hides while it significantly (p<0.05) increased the ash, protein and lipid contents. The derivable metabolic water and energy content calculation (table 4) showed that singeing significantly (p<0.05) significantly increases both the derivable metabolic water content and energy content of the hide.

**Table 1:** Heavy metal composition of the samples.

| **Parameter** | **Sample A** | **Sample B** | **Sample C** | **Mean** | **SD** | **%CV** |
| --- | --- | --- | --- | --- | --- | --- |
| (mg/kg) | (SWST) | (SWFW) | (control) | (X) |  |  |
| **Mercury (Hg)** | ND | ND | ND | - | - | - |
| **Lead (Pb)** | ND | ND | ND | - | - | - |
| **Cadmium (Cd)** | ND | ND | ND | - | - | - |
| **Cobalt (Co)** | ND | ND | ND | - | - | - |
| **Iron (Fe)** | 171.20<sup>a</sup> | 144.68<sup>b</sup> | 135.66<sup>b</sup> | 15.05 | 1.52 | 10.11 |
| **Copper (Cu)** | 0.75<sup>a</sup> | 0.60<sup>b</sup> | 2.67<sup>c</sup> | 0.13 | 0.09 | 70.30 |
| **Arsenic (As)** | ND | ND | ND | - | - | - |

Means with different superscripts (a, b, c) on the same row are significantly different at 5% ( $p < 0.05$ ). CV = coefficient of variation; SD = standard deviation; ND = not detected; SWST = singed with scrap tyre; SWFW = singed with firewood.

**Table 2:** Polycyclic aromatic hydrocarbon contents of samples.

| Parameter<br>(ppm) | Sample A<br>(SWST) | Sample B<br>(SWFW) | Sample C<br>(control) | Mean<br>(X) | SD | %CV |
| --- | --- | --- | --- | --- | --- | --- |
| Naphthalene | 5.63e<sup>-5a</sup> | 4.17e<sup>-6b</sup> | 3.5e<sup>-6c</sup> | 2.13e<sup>-5</sup> | 2.47e<sup>-5</sup> | 115.81 |
| 2-methyl Naphthalene | 2.17e<sup>-6a</sup> | 2.46e<sup>-6b</sup> | 9.82e<sup>-6c</sup> | 4.82e<sup>-6</sup> | 3.54e<sup>-6</sup> | 73.53 |
| Acenaphthylene | 3.63e<sup>-6a</sup> | 2.20e<sup>-6b</sup> | 3.44e<sup>-5c</sup> | 1.34e<sup>-5</sup> | 1.69e<sup>-5</sup> | 110.72 |
| Acenaphthene | 3.81e<sup>-6a</sup> | 2.57e<sup>-6b</sup> | 1.13e<sup>-4c</sup> | 3.98e<sup>-5</sup> | 5.17e<sup>-5</sup> | 130.08 |
| Flourene | 1.07e<sup>-5a</sup> | 1.13e<sup>-6b</sup> | 1.17e<sup>-4c</sup> | 4.29<sup>-5</sup> | 5.24e<sup>-5</sup> | 122.28 |
| Phenanthrene | 2.05e<sup>-5a</sup> | 1.17e<sup>-5b</sup> | 1.07e<sup>-4c</sup> | 4.63e<sup>-5</sup> | 4.29e<sup>-5</sup> | 92.62 |
| Anthracene | 5.53e<sup>-5a</sup> | 1.54e<sup>-6b</sup> | 1.04e<sup>-4c</sup> | 5.35e<sup>-5</sup> | 4.17e<sup>-5</sup> | 77.98 |
| Flouranthene | 3.55e<sup>-4a</sup> | 1.18e<sup>-4b</sup> | 2.60e<sup>-4c</sup> | 2.44e<sup>-4</sup> | 9.77e<sup>-5</sup> | 40.01 |
| Pyrene | 3.36e<sup>-4a</sup> | 1.93e<sup>-4b</sup> | 1.06e<sup>-4b</sup> | 2.12e<sup>-4</sup> | 9.50e<sup>-5</sup> | 44.87 |
| Benz[a]anthracene | 4.27e<sup>-4a</sup> | 1.85e<sup>-3b</sup> | 8.16e<sup>-4c</sup> | 1.03e<sup>-3</sup> | 5.99e<sup>-4</sup> | 58.18 |
| Chrysene | 5.61e<sup>-4a</sup> | 1.75e<sup>-4b</sup> | 8.68e<sup>-5c</sup> | 2.74e<sup>-4</sup> | 2.24e<sup>-4</sup> | 75.16 |
| Benzo[b]flouranthene | 1.24e<sup>-2a</sup> | 2.46e<sup>-3b</sup> | 1.76e<sup>-4c</sup> | 5.02e<sup>-3</sup> | 5.31e<sup>-3</sup> | 105.84 |
| Benzo[k]flouranthene | 2.33e<sup>-3a</sup> | 1.35e<sup>-4b</sup> | 9.50e<sup>-5c</sup> | 8.52e<sup>-4</sup> | 1.04e<sup>-3</sup> | 122.32 |
| Benzo[a]pyrene | 2.19e<sup>-3a</sup> | 2.30e<sup>-3a</sup> | 3.01e<sup>-4b</sup> | 1.60e<sup>-3</sup> | 9.19e<sup>-4</sup> | 57.49 |
| Indeno[1,2,3-cd]pyrene | 5.71e<sup>-5a</sup> | 1.06e<sup>-5b</sup> | 3.03e<sup>-4c</sup> | 1.24e<sup>-4</sup> | 1.28e<sup>-3</sup> | 103.85 |
| Dibenz[a,h]anthracene | 1.53e<sup>-4a</sup> | 3.22e<sup>-6b</sup> | 1.37e<sup>-3c</sup> | 5.08e<sup>-4</sup> | 6.12e<sup>-4</sup> | 120.33 |
| ΣPAH4 | 1.55e<sup>-2a</sup> | 6.79e<sup>-3b</sup> | 1.38e<sup>-3c</sup> | 7.89e<sup>-3</sup> | 3.37e<sup>-5</sup> | 699.73 |

Means with different superscripts (a, b, c) on the same row are significantly different at 5% (p<0.05). CV = coefficient of variation; SD = standard deviation; SWST = singed with scrap tyre; SWFW = singed with firewood; e<sup>-</sup> = exponential (×10<sup>-</sup>). 1 ppm = 1 mg/kg = 1000 µg/kg.

**Table 3:** Proximate composition of the samples.

| **Parameter** | **Sample A** | **Sample B** | **Sample C** | **Mean** | **SD** | **%CV** |
| --- | --- | --- | --- | --- | --- | --- |
| (%) | (SWST) | (SWFW) | (control) | (X) |  |  |
| **Moisture** | 52.47<sup>a</sup> | 55.36<sup>a</sup> | 62.19<sup>b</sup> | 56.67 | 4.12 | 7.27 |
| **Ash** | 1.03<sup>a</sup> | 1.02<sup>a</sup> | 0.28<sup>b</sup> | 0.78 | 0.35 | 45.25 |
| **A.CHO** | 0.21<sup>a</sup> | 0.21<sup>a</sup> | 0.85<sup>b</sup> | 0.78 | 0.30 | 71.71 |
| **Protein** | 44.98<sup>a</sup> | 41.17<sup>a</sup> | 34.49<sup>b</sup> | 40.21 | 0.30 | 10.79 |
| **Lipid** | 1.10<sup>a</sup> | 1.90<sup>b</sup> | 1.00<sup>a</sup> | 1.33 | 0.40 | 30.29 |
| **Fibre** | 0.21<sup>a</sup> | 0.34<sup>b</sup> | 1.19<sup>c</sup> | 0.58 | 0.43 | 74.61 |

Means with different superscripts (a, b, c) on the same row are significantly different at 5% ( $p < 0.05$ ). CV = coefficient of variation; SD = standard deviation; SWST = singed with scrap tyre; SWFW = singed with firewood; A.CHO = available carbohydrate.

**Table 4:**
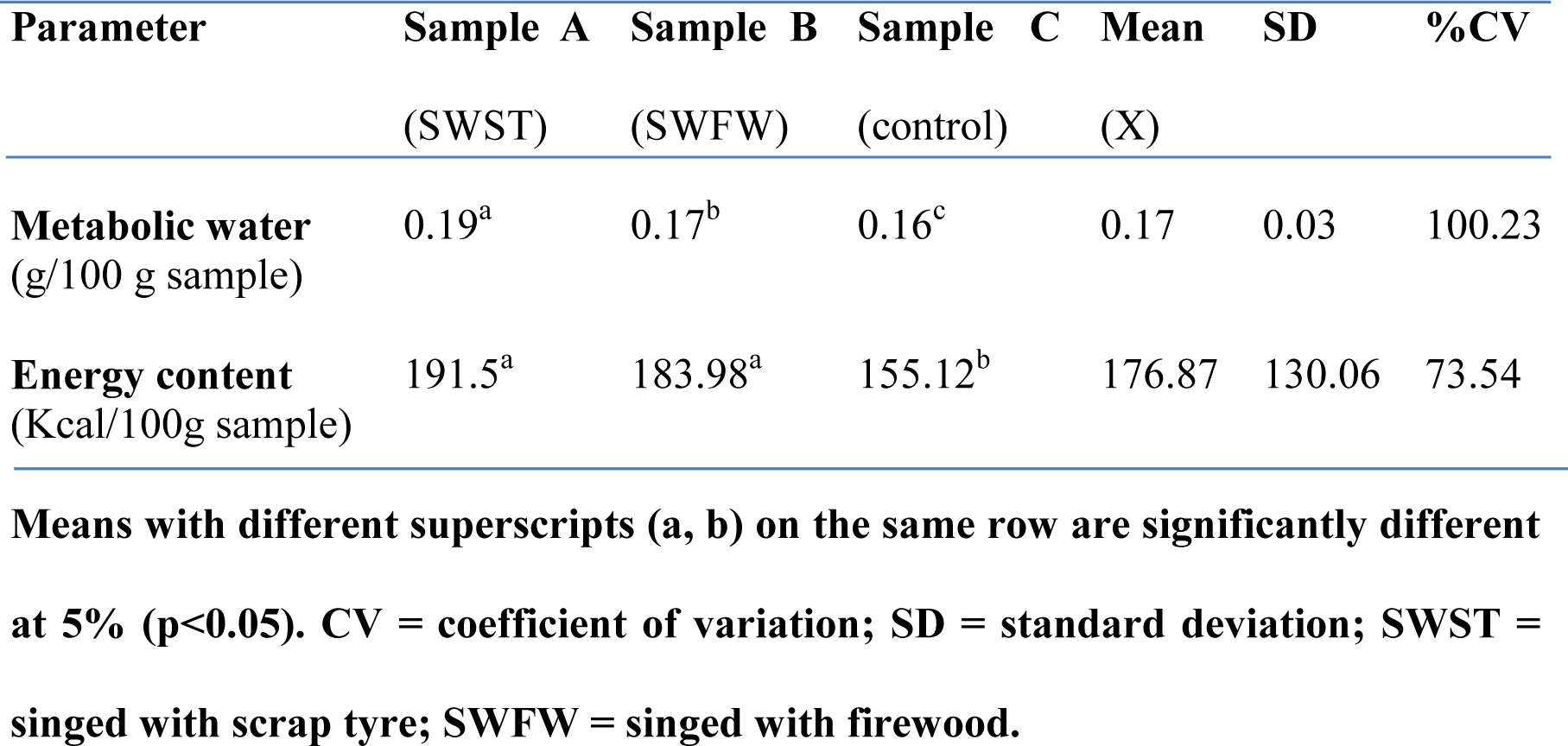
Metabolic water and energy content of the samples.

| **Parameter** | **Sample A** | **Sample B** | **Sample C** | **Mean** | **SD** | **%CV** |
| --- | --- | --- | --- | --- | --- | --- |
|  | (SWST) | (SWFW) | (control) | (X) |  |  |
| **Metabolic water**<br>(g/100 g sample) | 0.19<sup>a</sup> | 0.17<sup>b</sup> | 0.16<sup>c</sup> | 0.17 | 0.03 | 100.23 |
| **Energy content**<br>(Kcal/100g sample) | 191.5<sup>a</sup> | 183.98<sup>a</sup> | 155.12<sup>b</sup> | 176.87 | 130.06 | 73.54 |

**Means with different superscripts (a, b) on the same row are significantly different at 5% ( $p < 0.05$ ). CV = coefficient of variation; SD = standard deviation; SWST = singed with scrap tyre; SWFW = singed with firewood.**

## Discussion

Among the seven (7) heavy metals analyzed only iron and copper were detected. Sample A recorded the highest iron content and was significantly (p<0.05) different from the other samples. This indicates that singeing with scrap tyre increases the iron (Fe) concentration of hides. The significant (p<0.05) difference between sample A and sample B could be attributed to the presence of iron (Fe) in scrap tyres. The values recorded were below the average lethal dose for iron (Fe) which is reported to be in the range of 200 to 250 mg/kg. These notwithstanding, it was not certain whether the value obtained in the study posed any significant health implications to consumers because no maximum permissible level (MPLs) have been assigned to iron in animal foods including meat and fish according to MFPO (Meat Food Product Order), and some international standards like WHO and FDA including the Bulgarian standards (MFPO, 1973; Kamaruzzaman *et al*., 2010; Stancheva, 2010). The observed copper (Cu) levels in the 3 samples were far below the maximum permissible levels (MPLs) allowed in meats and meat products. The United States Department of Agriculture (USDA) reported the maximum permissible level of copper in meat as 20 mg/kg (USDA, 2006). The results of this study also suggest that processing hides by singeing and then washing significantly (p<0.05) reduces its copper (Cu) concentration, this could be observed with the two processed samples (A and B) when compared with the control (sample C). The significant (p<0.05) difference observed when samples A and B are compared could be attributed to the presence of copper in scrap tyres. Research reports indicate that singeing is not the only means by which animals could pick up heavy metal residues, but also, the soil, feed and drinking water are potential avenues from which these heavy metal residues could be picked up by the animals (Qui *et al*., 2008). This therefore explains why there were heavy metal residues in the unsigned (sample C) cattle hide.

However, the levels of heavy metals detected in all the hides in this study were generally quite low when compared to some cases of heavy metal concentrations in hides and other meat products reported by other researchers (Obiri-Danso *et al*., 2008; Okiei *et al*., 2009; Akwetey *et al*., 2013; Ijeoma *et al*., 2015).

The PAH concentrations in the various hide samples were analyzed to quantify the presence of USEPA 16 priority PAHs. All the 16 PAHs were detected at varying concentrations and significantly differed (p<0.05) in concentrations among the samples except benzo[a]pyrene which showed no significant difference between sample A and sample B. Benzo[b]flouranthene had the highest concentration among the PAHs analyzed and this occurred in sample A, followed by sample B. The PAH with the second highest concentration was Benzo[k]flouranthene in sample A while benzo[a]pyrene had the third highest concentration as recorded in sample B followed by sample A. with the exception of naphthalene, 2-methynaphthalene, acenaphthalene and indeno[1,2,3-cd]pyrene, all the analyzed PAHs recorded a concentration above 0.1 μg/kg in at least one of the samples. It was observed that processing cattle hides by singeing significantly increases the concentrations of some of the PAHs in the hides. The low concentration of some of the analyzed PAHs in the singed samples (sample A and B) compared to the control (sample C) could be attributed to the high level of deep scraping involved while washing the singed hides. It is reported that the concentration of PAH are generally on the surface than in the internal tissues. Consequently, careful washing might remove an average of up to 50% of the total PAHs (European commission, 2002). The concentration of chrysene (10.36 μg/kg, and 2.95 μg/kg), Benz[a]anthracene (12.7 μg/kg, and 7.23 μg/kg), Benzo[a]pyrene (8.76 μg/kg), Dibenz[a,h]anthracene (0.97 μg/kg), and Indeno[1,2,3-cd]pyrene (64 μg/kg) as reported by Manda for smoked meat and Amos-Tautua for *Suya* meat respectively was far higher than the results obtained in this study (Manda *et al*., 2012; Amos-Tautua *et al*., 2013). Only Benzo[b]flouranthene was lower compared to sample A in this study.

It has been estimated that nearly 70% of PAHs are consumed with food, including the consumption of smoked meat. Benzo[a]pyrene (B[a]P) for a long time was a marker of total PAHs content in food and environmental analysis. However, in many cases, B[a]P constitutes only 1–20% of the total PAHs content in the examined matrix. According to European Food Safety Authority (EFSA), benzo[a]pyrene is not a sufficient, suitable marker for indicating the presence of PAHs in food (EFSA, 2008). Thus, they assessed that the sum content of the four PAH compounds (“ΣPAH4”) called PAH4: Benzo[a]pyrene (B[a]P), Benzo[a]anthracene (B[a]A), Benzo[b]flouranthene (B[b]F) and Chrysene (CHR) is the most suitable indicator of PAHs in food. In 2011, based on the opinion of EFSA, modifications of the numerical limits of permissible PAH levels for individual food groups were carried out. Since September 1, 2014, the maximum level for the sum of PAH4 has been 12.00 μg/kg of a product (European Commission, 2002). The sum of PAH4 (“ΣPAH4”) in sample A in this study was higher than the EFSA recommended maximum level, while that of samples B and C was very low. Taking into account properties of several PAH compounds (i.e PAH4), the Scientific Committee on Food (SCF) recommended that the PAH contents in smoked meat products should be as low as reasonably achievable (Miculis, 2011).

The proximate compositions of these three (3) samples of cattle hide showed that Sample C had the highest moisture content which was significantly (p<0.05) higher than all other samples. Sample A had the least moisture content. The greater moisture content of sample C might be due to the fact that it did not pass through any process of heating during the course of its processing. While the lower moisture content observed with samples A and B might be attributed to the fact that heating was a common processing technique employed during the course of their preparation. Preparation of sample A involved singeing (burning in open fire) with scrap tyre as source of fuel, while that of sample B involved singeing with firewood as source of fuel. The moisture content obtained from the samples in this study was above the range (17.37% - 20.01%) of moisture contents of boiled and singed hides gotten from Ogbor hill abattoir reported by Ijeoma *et al*. (2015), and lower than that reported for beef (lean) by FAO (2015). Samples with high moisture content have low shelf life because micro-organisms thrive or grow more in food with high moisture content (Adepoju *et al*., 2006; Okonkwo and Opara, 2010).

The percentage ash content of the samples indicates that singeing significantly (p<0.05) increases the ash contents of the hides. This could be attributed to the heat intensity generated from the processing methods. Sample A and B recorded low values compared to what was reported by Akwetey, and Dabuo for hides processed by singeing with scrap tyre (1.50%) and firewood (0.83%), respectively, and also for beef (lean) reported by FAO (Dabuo, 2011; Akwetey *et al*., 2013; FAO, 2015). Sample C also recorded low value compared to what was reported for raw cow hides (0.87%) by Maduforo (2016). These low ash contents was indicative of low mineral content as opined by Francis (2007) that the ash content of any sample is a measure of likely mineral content of such a sample. The carbohydrate values significantly (p<0.05) decreased with singeing. The values for sample A and B was lower to the values (0.71%) recorded by Ijeoma *et al*. (2015) for tyre singed cow hides gotten from Ogbor hill abattoir.

The protein values significantly (p<0.05) increased with the singed samples (A and B) when compared with the control (sample C). Though there was no significant (p<0.05) difference between sample A (singed with scrap tyre) and sample B (singed with firewood), there was a significant increase between the control and the singed samples. Sample A had the highest protein value, and its values was far above the figures (8.68%) and (22.3%) reported by Ijeoma and FAO for tyre singed hides from Ogbor hill abattoir, and beef (lean) respectively (FAO, 2015; Ijeoma *et al*., 2015). Except sample B, all samples showed lower lipid content compared to what FAO (2015) reported for beef (lean) meat.

The fibre content of the three (3) samples (A, B and C) significantly (p<0.05) differed from each other. The control (sample C) had the highest fibre content, followed by sample B, and then sample A. From the results it is indicative that the extent of heating involved in processing the hide had a direct influence on the fibre content; the higher the heat the lower the fibre content. For example, the heat generated by scrap tyre while processing sample A is assumed to be higher than the heat generated by firewood while processing sample B, thus, the significant decrease in fibre content between sample B and A. The control (sample C) which did not pass through any heating process during the course of preparation had the highest fibre content. It has been reported that crude fibre content represents the amount of indigestible sugar present in a sample and is implicated in the maintenance of normal peristaltic movement of the intestinal tract. Hence, diets containing low fibre could cause constipation and eventually lead to disease of the colon like pile, appendicitis and cancer (Omosuli, 2014). The results obtained from calculating the derivable metabolic water content and energy content of the three hide samples indicated that singeing significantly increases both the derivable metabolic water and energy contents of the hides. There was significant difference among the three samples when their derivable metabolic water was compared.

## 4. CONCLUSION

Though singeing cattle hides with scrap tyre or firewood increases the energy content and derivable metabolic water of the hides, the practice introduces heavy metals and polycyclic aromatic hydrocarbons into the meat which makes it unhealthy for consumption and puts the unsuspecting consumer at health risk. It is therefore advised that singeing of animals (hide or any other part) be discouraged and more awareness created so that people will know the dangers to health posed by this harmful practice.

